# Reproductive state controls transcription in the murine liver, with implications for breast cancer liver metastasis

**DOI:** 10.1101/2024.08.02.606434

**Authors:** Michelle K. Ozaki, Yi Zhang, Alexandra Q. Bartlett, Elise de Wilde, Xiangnan Guan, Alex Yang, Zheng Xia, Pepper Schedin

## Abstract

Liver biology is functionally linked to lactation, as liver size and metabolic output increase during lactation to support synthesis of breast milk. Upon weaning, the rodent liver returns to baseline homeostasis via hepatocyte cell death, in a process considered liver involution. To explore liver biology changes across a lactation-wean cycle, we employed transcriptomic profiling. We identified elevated hepatocyte proliferation and anabolic metabolism gene signatures during lactation, consistent with the liver being a major producer of substrates needed for milk production. Rapid loss of these capacities upon weaning correlated with catabolic metabolism, lysosomal-mediated cell death, and an influx of immune suppressive cells. Furthermore, we identified that the transcriptional profiles associated with liver involution share similarities with the gene expression patterns of liver pre-metastatic niches. This work identifies features of reproductive control of liver biology that sets a foundation for better understanding the potential role of the liver in maternal health.

## Introduction

The liver plays a crucial role in maintaining healthy blood nutrient levels by regulating nutrient uptake, storage, and release into the circulatory system. To accommodate distinct metabolic demands due to differences in adult body size, a mechanism referred to as the hepatostat^1^ maintains direct proportionality between liver and body size. As such, in healthy adult individuals with stable weight, liver size does not fluctuate. Rather, changes in adult liver size are frequently associated with liver pathologies such as cirrhosis^2^, fatty liver disease^3,4^ and cancer^5^. However, recent studies in rodents have found that the liver undergoes pregnancy and lactation induced growth, followed by weaning-induced involution^6–8^, similar to that which occurs in the mammary gland^9–11^. Evidence for human relevance is suggested by recent observations in healthy women. Similar to rodents, the human liver undergoes increases and decreases in size over the course of a reproductive cycle that is also independent of body mass^7^. Little is known about the molecular pathways associated with liver remodeling that occurs with lactation and weaning. Also unknown is how these molecular changes may affect sex-specific liver function or pathologies. Thus, understanding these mechanisms is fundamental to our understanding of liver biology in women. For example, tissue growth followed by involution can occur through cycles of cellular hypertrophy and hypotrophy^12^, or by cellular proliferation followed by programmed cell death^11^. With respect to physiology, the former is thought to be immune silent, whereas the latter can trigger immune regulatory responses^13^, which can have implications on organ function and health^14,15^.

Cell death during weaning-induced mammary gland involution is accompanied by classic wound-healing responses such as activation of fibroblasts^16^, and increased ECM deposition^17,18^, immune cell infiltration^19,20^, and active immune suppression^21–23^. In the liver, similar stromal changes are known to associate with several liver pathologies including liver cirrhosis^24^, fatty liver disease^25,26^, autoimmune hepatitis^27^, and cancer^28,29^. Of potential relevance to cancer, weaning-induced involution of the breast and liver is associated with breast cancer promotion and increased liver metastasis in both rodent models^6,8^ and women^30–32^. In sum, these prior findings implicate postpartum liver remodeling as a potential mediator of liver pathologies in recently pregnant women. Here, we deeply characterize the transcriptional profiles in the murine liver across a lactation-wean cycle and report the underlying molecular changes associated with these distinct reproductive states. Our goal is to elucidate previously underrecognized liver biology and provide molecular insight into reproductive driven liver physiology and pathology relevant to women.

## Results

### Transcriptional profiling of the murine liver reveals distinct reproductive states

The mammary gland undergoes dramatic functional changes as it transitions from a non-pregnant (nulliparous) state to lactation, and again upon weaning to an involuted, non-secretory state. These changes can be observed at the histologic level (Fig 1A upper panels). As such, the rodent mammary gland is a useful model to investigate how gene expression changes impact differentiation states and tissue remodeling processes. However, unlike the mammary gland, gross morphologic and histologic characteristics of the liver do not markedly change across a lactation-wean cycle in rodents (Fig 1A, bottom panels), even though reproductive control of liver size and function have been described at the cell and protein levels^6–8,33^.

**Fig 1.**
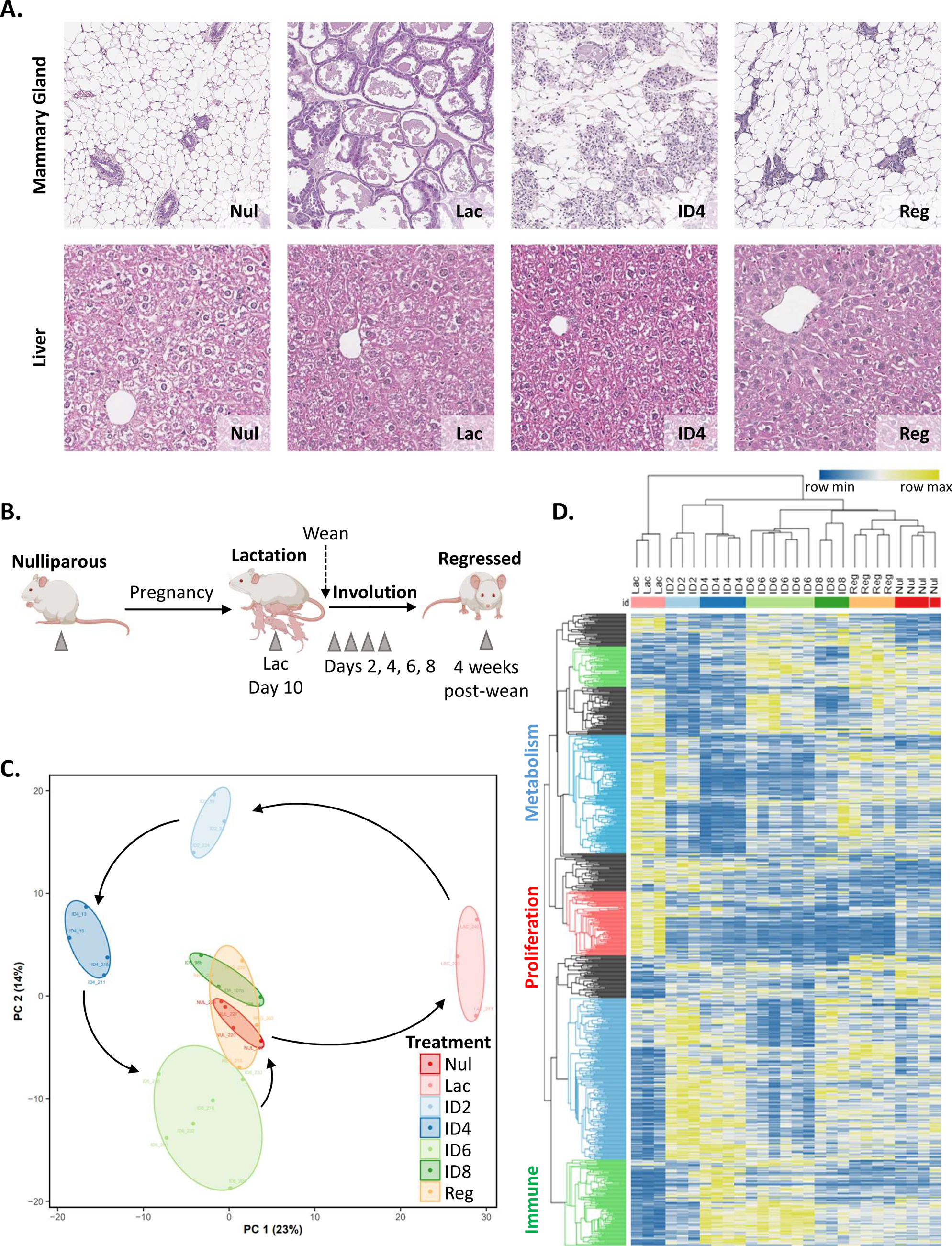
Liver transcriptional profiles differ by reproductive state. **A)** Hematoxylin & eosin (H&E) staining of mammary glands (top) and livers (bottom) from nulliparous (Nul), lactation (Lac), involution day 4 (ID4), and 28 days post wean/regressed (Reg) mice. **B)** Study outline for collecting livers at specific reproductive states depicted by grey triangles for bulk RNAseq. **C)** Principal component (PC) analysis of the top 5000 most variable genes between reproductive states. **D)** Unsupervised hierarchal clustering of the top 100 most differentially expressed genes between each reproductive group (841 genes total, lowly expressed-blue, highly expressed-yellow). Animals are clustered horizontally, and genes vertically, using one-minus kendall’s correlation. Families of related genes fell into three main categories: proliferation (red), metabolism (blue), immune (green), with unrelated clusters in black.

To more extensively characterize the alterations in liver biology during a lactation-wean cycle, we performed bulk tissue RNA-sequencing on mouse livers from nulliparous (baseline), lactation day 10 (full lactation state), post-wean days 2, 4, 6, 8 (the involution window), and fully regressed/parous (4 weeks post-wean) mice (Fig 1B). Principal component (PC) analysis conducted on the top 5000 most variably expressed genes across all samples demonstrated distinct gene expression profiles associate with each reproductive state (Fig 1C). These data support the hypothesis that discrete functional states of the liver occur across a reproductive cycle despite the lack of obvious changes to liver morphology. In addition to PC analysis, we conducted hierarchal clustering, which similarly revealed sample clustering by reproductive state. Hierarchal clustering of genes (y-axis) identified 3 major clusters of functionally related genes that differed between reproductive groups (Fig 1D): proliferation (red), metabolism (blue), and immune (green), as well as several clusters of genes that did not cluster into known biologic pathways (black). Combined, these data revealed lactation group samples had the most distinct gene expression pattern from all the other groups (Fig 1C, D). Further, nulliparous, involution day 8, and regressed samples largely overlapped. This likely reflects shared liver function and suggests that involution is largely complete by involution day 8. Involution days 2, 4, and 6 all clustered distinctly from all other groups and from each other (Fig 1C, D), consistent with the involution process being dynamic and rapid. Collectively, these data highlight how liver gene expression reflects the uniquely tuned interplay between liver function and reproductive state.

### The lactation-state liver is characterized by increased cell proliferation and anabolic metabolism

To assess how liver gene expression might change to support the demands of lactation, we performed two group comparisons between the nulliparous and lactation groups. We expanded our analysis to the top 500 DEGs to better leverage the depth of RNA-Seq data. To identify the main biological pathways that are differentially regulated, we employed STRING analysis (the Search Tool for the Retrieval of Interacting Genes/Proteins) (Fig 2A). STRING coupled with gene ontology (GO) enrichment analysis^34,35^ identified 3 main gene clusters that differed between nulliparous and lactation states. These pathways were associated with the biological functions of proliferation (red), metabolism (blue) and immunity (green) (Fig 2B, Supplementary Figure 1A-B). Next, we used gene set enrichment analysis (GSEA) to determine which reproductive group corresponded with elevated proliferation. Proliferation genes associated with the hallmark G2M checkpoint pathway were upregulated ∼3-fold in lactation compared to the nulliparous group (Fig 2C). To see how these differences compared across a lactation-wean cycle, we next employed the use of single sample gene set enrichment analysis (ssGSEA)^36^. This analysis showed the Hallmark G2M pathway was significantly increased during lactation as compared to all other reproductive time points (Fig 2D). We next used ssGSEA to look at 10 independent proliferation gene signatures within the Molecular Signatures Data Base (MSigDB)^37^, and all 10 signatures showed enriched proliferation during lactation (Fig 2E). Of note, we found proliferation pathways were lowest in the regressed, parous liver, suggesting that the parous liver may have reduced proliferation compared to the nulliparous liver (Fig 2D, E, Supplementary Fig 1C-D). Next, we performed regulon analysis, which quantifies inferred transcription factor network activity rather than specific gene regulation. Regulon analysis identified transcription factors associated with proliferation (e.g. CUX1, MYC, E2F1) upregulated in the lactation group (red) compared to the nulliparous group (blue) (Fig 2F, green). In sum, these data show increased proliferation is a key attribute of the liver during lactation. These data correlate well with the reported increase in liver size in humans^7^ and mice^6^ during pregnancy and lactation.

**Fig 2.**
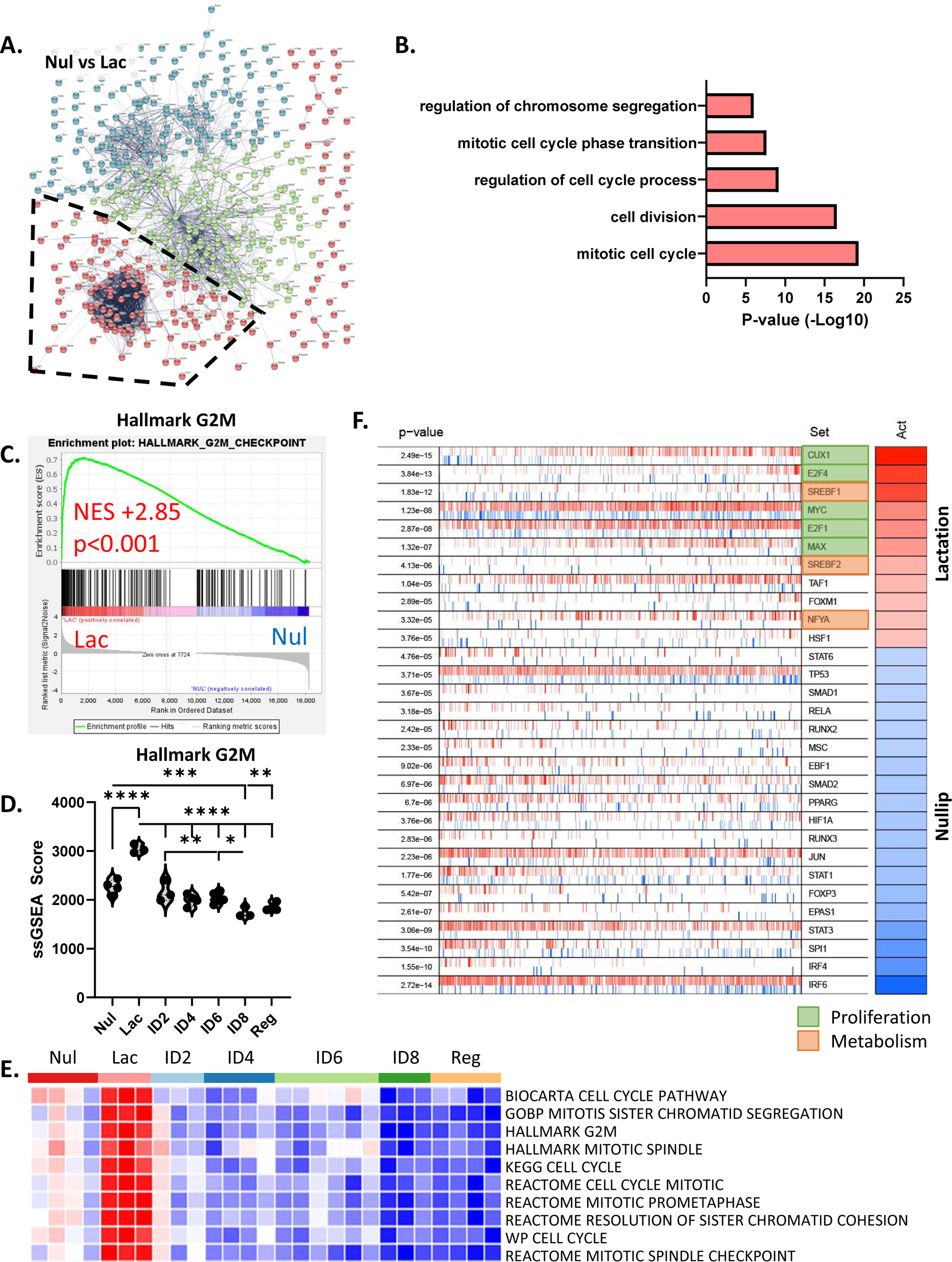
Lactation is a state of increased proliferation in the liver. **A)** Comparison of the top 500 differentially expressed genes nulliparous and lactation livers by STRING analysis, sorted by k-mean clustering into 3 main clusters of related genes. **B)** GO Term Enrichment of the top 5 families of genes in the red STRING cluster comparing nulliparous to lactation samples identified proliferation genes. **C)** GSEA of the Hallmark G2M checkpoint pathway comparing lactation liver samples compared to nulliparous liver samples. **D)** ssGSEA scores for the Hallmark G2M pathway for all reproductive groups (ANOVA, *p<0.05, **p<0.01, ***p<0.001, ****p<0.0001). **E)** Heatmap of ssGSEA scores run on publicly available pathways associated with cell proliferation pulled from the MSigDB. Values were normalized within each pathway (row). Increased pathway expression is displayed in red and decreased pathway expression in blue. **F)** Transcription factor regulon analysis between nulliparous (blue) and lactation (red) groups. Regulons associated with proliferation are highlighted in green and regulons associated with metabolism are highlighted in orange.

To further understand the unique gene expression attributes of the liver during lactation, we investigated genes associated with the metabolism cluster (blue cluster, Fig 3A). We evaluated gene signatures specific for anabolic metabolism pathways, as we hypothesized increased anabolic metabolites are required for the metabolically demanding process of milk production. First, we evaluated gene signatures for fatty acid and lipid metabolism, as lipids are a primary component of breast milk^38^. Using GSEA analysis, fatty acid synthesis was enriched over 2-fold during lactation compared to baseline nulliparous livers (Fig 3C). These data aligned with our regulon analysis (Fig2A, orange), which showed an enrichment in lipid metabolism regulons (e.g. SREBF1, SREBF2, NFYA) during lactation. We also found evidence for increased energy production in the liver during lactation as evidenced by increased expression of classical energy production pathways: glycolysis, Kreb cycle, and electron transport chain (ETC) (Fig 3C). To extend these observations across the reproductive cycle, we used the classical energy production gene pathways curated by the MSigDB to perform ssGSEA analysis with multiple group comparisons across all time points. From these analyses, we confirmed fatty acid synthesis pathways were highest at lactation compared to any other time point (Fig3D). Glycolysis, Kreb cycle, and ETC pathways also peaked during lactation. Overall, our data indicate that lactation is a period of increased metabolic demand, increased lipid synthesis, and increased overall energy production for the liver. These findings are consistent with the liver having a key role in mammary milk production (Fig 3F).

**Fig 3.**
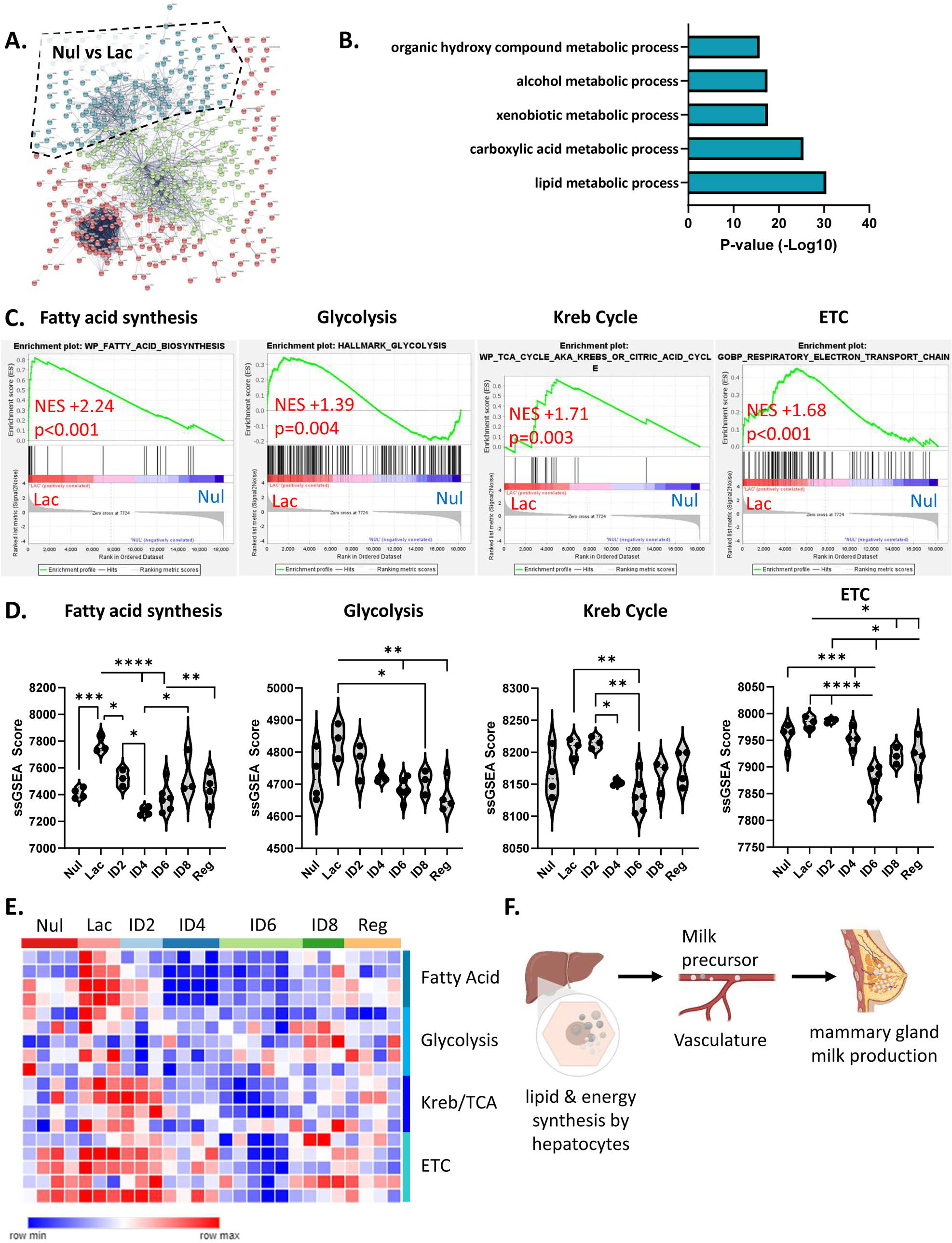
Lactation is a state of increased liver energy production. **A & B)** Analysis of the blue STRING cluster by GO Term Enrichment. Analysis the top 5 families of related genes between nulliparous and lactation livers. **C)** GSEA analysis comparing nulliparous livers and Iactation group livers for energy production pathways: fatty acid synthesis, glycolysis, Kreb cycle, and electron transport chain (ETC) pathways. **D)** ssGSEA scores for fatty acid synthesis, glycolysis, Kreb cycle, and ETC pathways for all reproductive groups (one-way ANOVA, *p<0.05, **p<0.01, ***p<0.001, ****p<0.0001). **E)** Heatmap of ssGSEA values run on publicly available pathways associated with glycolysis, Kreb cycle, electron transport chain, or fatty acid synthesis pulled from the MsigDB. Values are normalized within each pathway (row). Increased pathway expression is displayed in red and decreased pathway expression in blue. **F)** Model depicting liver production of substrates needed for mammary gland milk production.

### Involution is a state characterized by catabolic metabolism

In our metabolic pathway analyses across all groups, we identified a stark drop in anabolic pathways immediately post-wean (Fig 3D). To capture early shifts in metabolism following wean, we compared lactation to our earliest involution timeout, day 2. STRING analysis identified two primary clusters of genes that differed between lactation and involution day 2. These were changes in proliferation (red), as previously described above (Fig 2A, B), and catabolic metabolism (green) (Fig 4A, B). Using GSEA analysis, we investigated gene expression profiles of specific catabolic pathways including: amino acids, lipids, sugars, and nucleic acids. GSEA analysis identified amino acid catabolism as significantly enriched during early involution (Fig 4C-D). In addition to amino acid catabolism, we identified a strong enrichment for lipid catabolism pathways during involution day 2 (Fig 4E). Peak lipid catabolism occurred at involution day 4 as assessed by ssGSEA analysis (Fig 4F). We validated these increases in amino acid and lipid catabolism by assessing multiple MSigDB pathways by ssGSEA (Fig 4G). Further, we looked at the catabolism of DNA nucleotides, mRNA nucleotides, and polysaccharides. We found these pathways were not significantly enriched during involution compared to lactation (Supplementary Fig 1E-G). Of note, halting milk synthesis products (e.g. lipid synthesis) does not necessarily imply a need to breakdown metabolites. Thus, the observed increase in catabolism could suggest that the liver is responding to milk stasis that occurs upon weaning by supporting the breakdown of excess circulating milk precursors such as lipid (Fig 4G).

**Fig 4.**
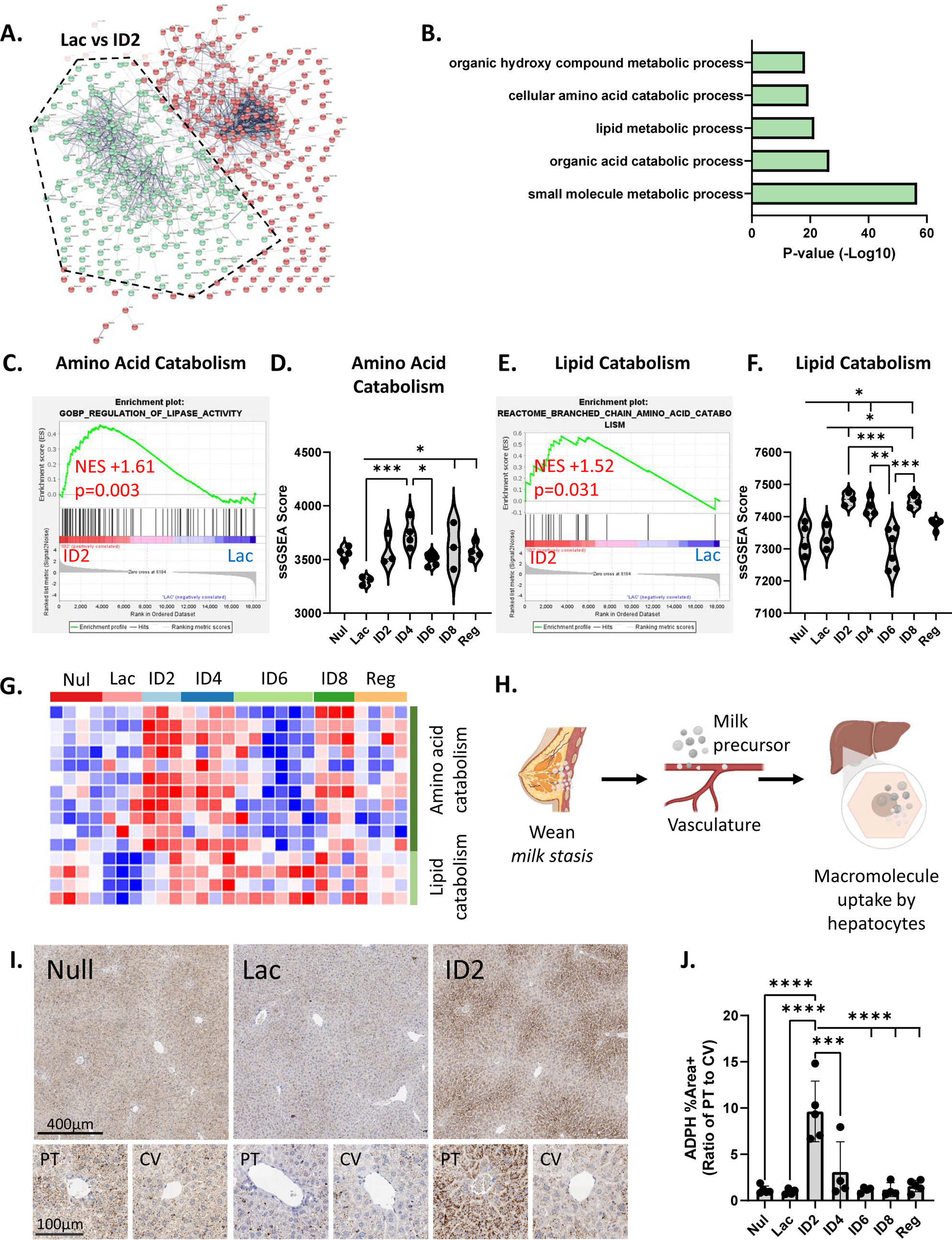
Weaning associates with catabolic breakdown of macromolecules by the liver. **A)** Comparison of the top 500 differentially expressed genes between lactation and involution day 2 livers by STRING analysis, sorted by k-mean clustering into 2 main clusters of related genes. **B)** GO Term Enrichment of the top 5 families of genes in the green STRING cluster comparing lactation to involution day 2 samples identified catabolic pathways. **C)** GSEA identified an enrichment of genes associated with amino acid catabolism in involution day 2 livers compared to lactation livers. **D)** ssGSEA scores of the amino acid catabolism pathway for all reproductive groups (ANOVA, *p<0.05, **p<0.01, ***p<0.001). **E)** GSEA analysis of lipid catabolism pathway genes between lactation and involution day 2 groups. **F)** ssGSEA scores for the lipid catabolism pathway across all groups using (ANOVA, *p<0.05, **p<0.01, ***p<0.001). **G)** Heatmap of ssGSEA scores run on additional pathways associated with amino acid catabolism (top, dark green) or lipid catabolism (bottom, light green). **H)** Model depicting milk stasis leading to buildup of milk precursor metabolites in the blood and uptake by the liver. **I)** Representative images of adipophilin stained by immunohistochemistry on nulliparous, lactation, and involution day 2 livers. Insets depict staining around portal triads (PT) regions compared to central vein (CV) regions. **J)** Quantification of adipophilin staining as a ratio of staining around portal triads compared to central veins (n=3-5 mice/group, one-way ANOVA, ***p<0.001, ****p<0.0001).

To further validate upregulation of the lipid catabolism during involution, we performed immunohistochemical staining on liver tissues across a reproductive cycle for the lipid binding protein adipophilin (ADPH)^39,40^. If the liver is responsible for lipid breakdown in response to milk stasis, we anticipated accumulation of lipid within the liver during involution, even though lipid synthetic pathways are downregulated (Fig 3D). Further, if lipid was coming in from circulation, potentially due to milk stasis, we anticipated lipid to be located near vasculature. We found ADPH protein levels were lowest during lactation (Fig 4I-J), supporting the idea that lipid is not stored in the liver during lactation, but rather rapidly secreted to support milk synthesis. Further, ADPH staining increased post-weaning (involution day 2), and interestingly, all ADPH staining was in hepatocytes located near portal triads, with very little staining near central veins (Fig 4I-J). This is consistent with lipids coming into the liver via the blood rather than new synthesis of lipid. In sum, these data support a working model whereby milk stasis in the mammary gland causes accumulation of milk-precursor macromolecules in the blood. This is followed by enhanced lipid and amino acid uptake in the liver, where these components are subsequently catabolized (Fig 4G).

### Hepatocyte cell death correlates with lysosomal lipid breakdown

Interestingly, excess lipid accumulation has been linked to cell death mechanisms in the involuting mammary gland^41^. In this type of cell death, lipid is shuttled into the lysosomes of epithelial cells, causing lysosomal instability and release of lysosomal contents, including lysosomal proteases^41,42^. The release of these proteases and stored lipid triggers cellular damage and caspase-mediated cell death^42^. Given our evidence for influx of lipid in the involuting liver, we hypothesized that excess lipid from milk stasis may be responsible for triggering cell death of hepatocytes in the involuting liver. To investigate this hypothesis, we first sought to validate previous reports of hepatocyte cell death during involution in our dataset^6^. We identified cell death pathways between involution day 2 to day 4 using STRING and PANTHER analyses (Fig 5A, B). By GSEA, apoptotic signatures were enriched during involution day 4 compared to involution day 2 (Fig 5C). Further, cell death pathways were at their lowest during lactation, with peak levels occurring during involution days 4 and 6 (Fig 5D). Next, we looked at 5 different apoptotic signatures from MSigDB by ssGSEA and confirmed that involution days 4 and 6 were enriched for all 5 apoptotic signatures (Fig 5D,E). We looked at enrichment of other cell death mechanisms, such as necrosis and autophagy, and found apoptotic signatures were the dominant enriched pathways (Supplementary Table 1).

**Fig 5.**
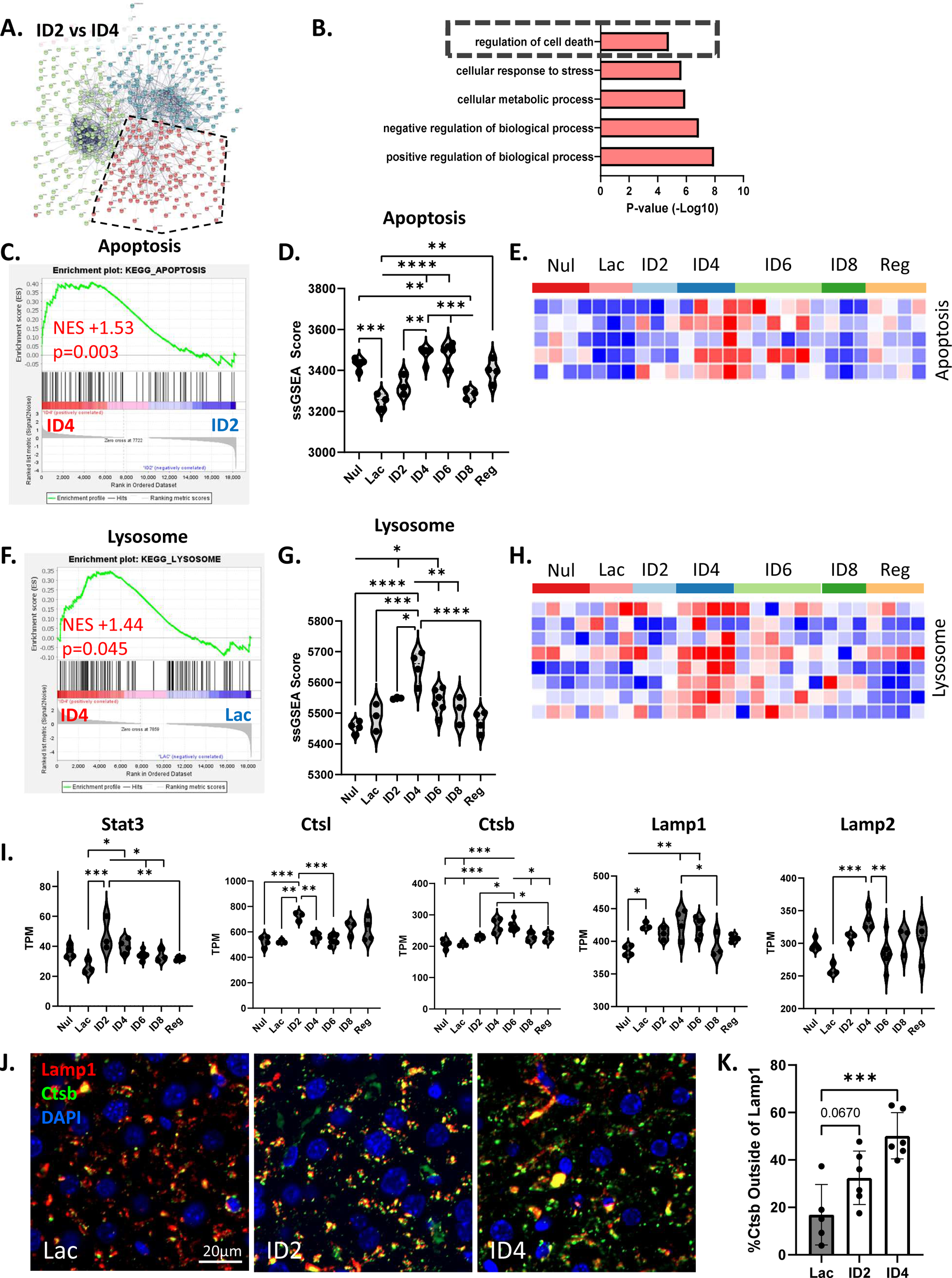
Cell death signature enrichment during liver involution correlates with lysosomal lipid breakdown. **A)** Comparison of the top 500 differentially expressed genes between involution day 2 and day 4 livers by STRING analysis, sorted by k-mean clustering into 3 main clusters of related genes. **B)** GO Term Enrichment of the top 5 families of genes in the red STRING cluster comparing involution day 2 to involution day 4 samples. **C)** GSEA analysis of the KEGG apoptosis pathway between involution day 4 livers compared to lactation group livers. **D)** ssGSEA scores for the KEGG apoptosis pathway assessed across all groups (ANOVA, **p<0.01, ***p<0.001, ****p<0.0001). **E)** Heatmap of ssGSEA scores for various apoptosis pathways from the MSigDB. **F)** GSEA analysis of the KEGG lysosomal compared between involution day 4 livers and lactation group livers. **G)** ssGSEA scores for the KEGG lysosome pathway were assessed across all groups by (one-way ANOVA, *p<0.05, **p<0.01, ***p<0.001, ****p<0.0001). **H)** Heatmap of ssGSEA scores for various lysosome pathways from the MSigDB. **I)** Transcripts per million for Stat3, cathepsin L (Ctsl) and cathepsin B (Ctsb), and lysosomal membrane proteins Lamp1 and Lamp2 across reproductive time points. **J)** Immunofluorescent staining to assess cathepsin B (green) localization within or outside of lysosomes (Lamp1, red) during lactation, involution day 2, and involution day 4. Scale bar represents 20µm. **K)** Quantification of cytoplasmic cathepsin B (one-way ANOVA, ***p<0.001).

To further investigate our hypothesis of lipid induced hepatocyte cell death, we next looked at whether gene expression during involution was consistent with increased lysosomal activity, as reported ^42^. We found lysosomal pathways to be significantly enriched in the liver during involution day 4 (Fig 5F). Of note, the lysosome is a major center for amino acid and lipid catabolism, and these findings were consistent with the concurrent enrichment of amino acid and lipid catabolism signatures we observed post-wean (Fig 4C-F). Lysosomal mediated cell death has been reported to be downstream of Stat3 activation^41–43^, and we identified an increase in *Stat3* expression at involution day 2 (Fig 5I). Transcription factor regulon and GSEA analysis further confirmed activation of the Stat3 pathway (Supplementary Fig 2A-B). Next, we looked at transcript levels for lysosomal structural proteins *Lamp1* and *Lamp2*, and lysosomal proteases, cathepsin L *(Ctsl)* and cathepsin b *(Ctsb)*, as increases in these transcripts associate with lysosomal membrane leakiness and cell death^42^. *Ctsl* expression was significantly enriched at involution day 2, while *Ctsb* expression peaked at involution days 4 and 6 (Fig 5I). Next, we evaluated expression of *Pmca2*, a negative regulator of lysosome mediated cell death^44^, and found *Pmca2* expression decreased during involution (Supplementary Figure 2C). In sum, these data are consistent with the hypothesis that lipid uptake following milk stasis may be an early event that triggers lysosomal membrane permeabilization, resulting in lysosomal mediated apoptosis.

To further investigate lysosomal mediated cell death, we used immunofluorescence to stain for Lamp1 (red), and Ctsb (green) (Fig 5J). Under non-apoptotic conditions cathepsins (Ctsb) should co-localize with lysosomes (Lamp1), whereas during lysosomal mediated cell death cathepsins will be found in the cytoplasm^41,42,44^. During lactation, Ctsb was predominantly located within lysosomes, with less than 20% of Ctsb located within the cytoplasm (Fig 5J-K). However, cytoplasmic Ctsb staining increased during involution day 2 and further increased at involution day 4, where ∼50% of total Ctsb was located in the cytoplasm (Fig 5J-K). These data are consistent with a model whereby increased lipid accumulation observed in hepatocytes shortly after weaning contributes to lysosomal membrane instability, release of cathepsin b, and subsequent lysosomal mediated apoptotic cell death of hepatocytes.

### Increase in immune suppressive myeloid cell phenotypes during involution

To further investigate liver involution, we investigated the blue cluster of genes that differ between involution days 2 and 4, and identified enrichment of immune signatures by STRING and GO Enrichment analyses (Fig 5A, 6A). GSEA analysis shows 2.2-fold enrichment of general inflammation associated genes (Hallmark Inflammatory Response) at involution day 4 compared to day 2 (Fig 6B). Further, using ssGSEA, we identified a decrease in inflammation genes during lactation, and highest enrichment of inflammation genes at involution day 4 (Fig 6C). Generally, these results indicate liver involution as a period of inflammation. To better characterize immune cell subtypes that might contribute to this inflammation, we employed CibersortX, which estimates relative immune cell compositions^45,46^. CibersortX identified monocytes as the main cell type present in the liver across all reproductive time points (Supplementary Fig 3A). When delineated by reproductive time point, monocyte signatures were most abundant during involution days 4 and 6 (Fig 6D). Next, we used multiplex immunohistochemistry (mIHC) staining to identify monocytes and their abundance in livers from different reproductive stages (Fig 6E). Similar to the CibersortX analysis, we identified monocytes (CD45+ CD11b+ Gr1+) cells were most abundant at involution day 4, and lowest at lactation when analyzing across multiple regions within the liver (Fig 6E-F, n=5-6 mice/group, n=2-3 ROIs/mouse). When averaged per mouse, data was consistent, with involution day 4 having the most CD45+ CD11b+Gr1+ cells (Supplementary Fig 3B).

**Fig 6.**
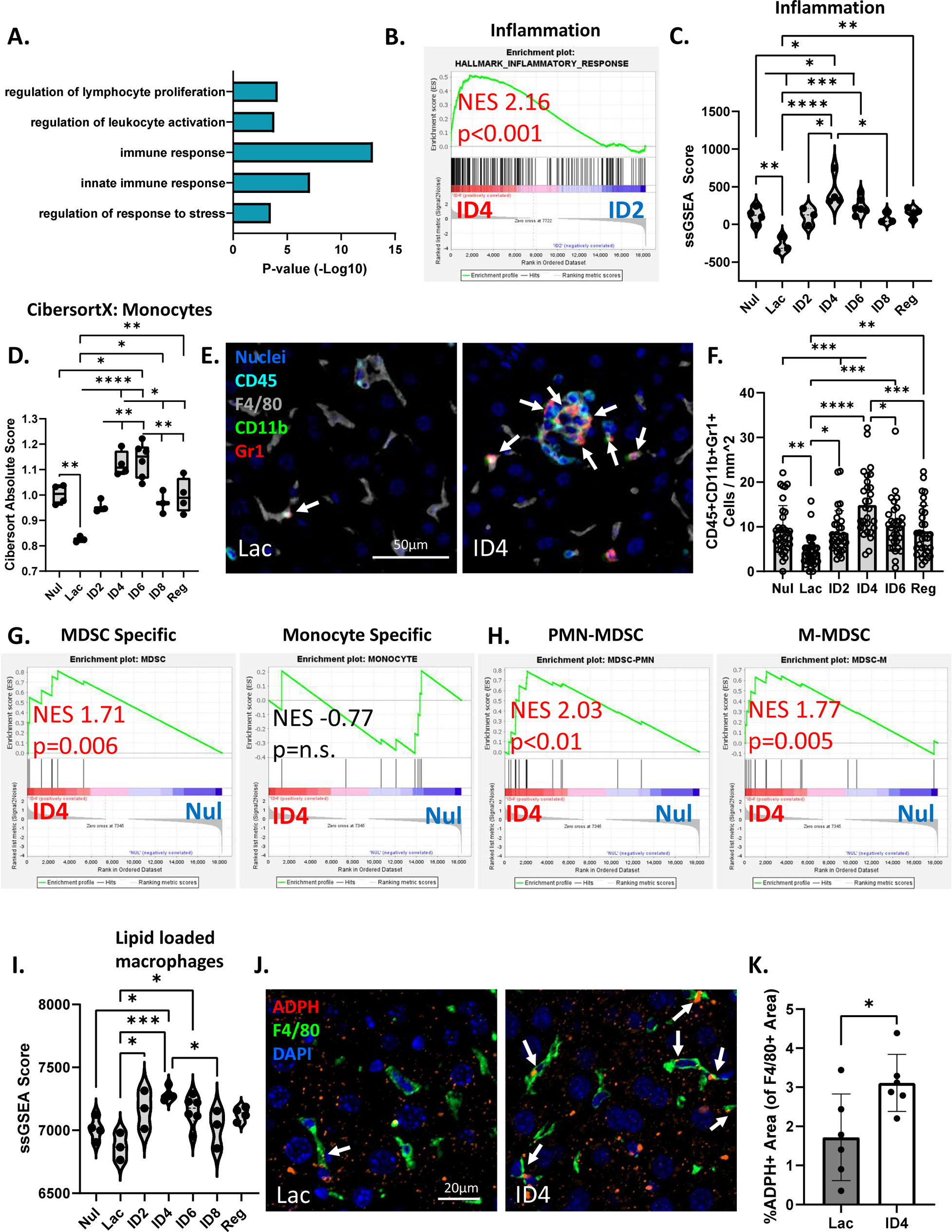
Immune cells present during liver involution are associated with immune suppression. **A)** GO Term Enrichment of the top 5 families of genes in the blue STRING cluster comparing lactation to involution day 2 samples. **B)** GSEA analysis of the Hallmark inflammatory response pathway between involution day 4 and involution day 2 livers. **C)** ssGSEA scores for the Hallmark inflammatory response pathway assessed across all groups by (ANOVA, *p<0.05, **p<0.01, ***p<0.001, ****p<0.0001). **D)** CibersortX analysis of inferred monocyte abundance at each reproductive time point. **E)** Pseudo-colored multiplex IHC images depicting CD45+ CD11b+ Gr1+ myeloid cells (white arrows) in lactation and involution day 4 livers. **F)** Quantification of CD45+CD11b+Gr1+ myeloid cells (n=6 mice/group, 4 regions per liver, one-way ANOVA, *p<0.05, **p<0.01, ***p<0.001), ****p<0.0001). **G)** GSEA analysis of MDSC-specific genes or monocyte-specific genes compared between involution day 4 livers and nulliparous group livers. **H)** GSEA analysis of PMN-MDSC and M-MDSC signatures between nulliparous and involution day 4 livers. **I)** Lipid loaded macrophage pathway genes were assessed across all groups by ssGSEA analysis (ANOVA, *p<0.05, **p<0.01, ***p<0.001). **J)** Liver sections IF stained for lipid (ADPH) and macrophages (f4/80) and **K)** dual staining quantified (n=6/group, student’s t-test, *p<0.05).

Given previous findings that identify liver involution as a period of increased breast cancer metastasis to the liver, which is mediated in part by transient immune suppression^8^, we assessed if the CD45+CD11b+Gr1+ myeloid cells present during involution day 4 exhibited immune gene signatures associated with myeloid derived suppressor cells (MDSCs). To address this, we first used gene signatures derived from Alshetaiwi, et al. that distinguish between MDSCs and monocytes^47^. To better understand how the immune milieu of involution differs from that of the normal healthy liver we compared involution day 4 livers to nulliparous. We found that signatures specific to MSDCs were enriched in our involution day 4 group compared to nulliparous (Fig 6G, left), while the signature specific to monocytes did not differ between these two groups (Fig 6G, right). MDSCs have been reported to originate from either a monocyte lineage, or a granulocytic lineage that can share features with neutrophils^48^. To address if our MDSCs could be neutrophils, we also used GSEA analysis to identify if a signature specific to neutrophils differed between nulliparous and involution day 4 livers. We found that neutrophil signatures were also not enriched (Supplementary Fig 3C). GSEA analysis on additional MDSC signatures^48^ revealed subtypes for both granulocytic (PMN-MDSCs) and monocytic (M-MDSCs) were enriched at involution day 4 compared to nulliparous (Fig 6H). These data suggest that the CD45+CD11b+Gr1+ myeloid cells detected by mIHC may be associated with both granulocytic and monocytic MDSCs.

Next, we wanted to investigate the presence of other myeloid cell subtypes and investigated M1 vs M2 macrophages. CibersortX revealed barely any anti-inflammatory M2 macrophage gene signatures, and no differences in M1 macrophages gene signatures over time (Supplementary Fig 3A, D). While we did not see differences in M1 and M2 macrophages by reproductive state, we sought to further investigate other subtypes of macrophages given their abundance in the liver^49^. Given the influx of lipid observed during involution, we next sought to investigate alterations in lipid loaded macrophages, which have been reported to have immune suppressive function^50–52^. We used a gene signature derived from lipid loaded macrophages obtained from breast, colon, and prostate cancer patients and mouse models^50^, and ran ssGSEA analysis (**Fig 6I**). This lipid loaded macrophage signature was most significantly enriched in involution day 4 livers and lowest in lactation livers (**Fig 6I**). GSEA analyses confirmed this lipid loaded macrophage signature was enriched at involution day 4, but not at involution day 6 (Supplementary Fig 3E). Next, we sought to validate the presence of lipid loaded macrophages during involution by staining for adipophilin positive F4/80+ macrophages by immunofluorescence (**Fig 6J**). Supporting our ssGSEA analysis, we identified an increase in adipophilin-positive macrophages during involution day 4 as compared to lactation group livers (**Fig 6J-K**).

### Involution has molecular attributes of a pre-metastatic niche, but not other pathological liver states

We anticipated the dramatic changes observed in the involuting liver, including catabolic metabolism, lipid mediated cell death, and immunosuppressive inflammation could have implications for liver pathologies. Here we investigate if involution resembles pathological liver states. Given previous findings that identified murine liver involution as a window of increased risk for breast cancer metastasis ^6,8^, we first looked for evidence for the establishment of a pre-metastatic niche during involution compared to nulliparous. We identified two gene signatures that describe the pre-metastatic niche in the liver, one generated from mouse and human studies on pancreatic and colon cancer^53^, and the other from a cohort of gastric cancer patients^54^. We identified significant enrichment in both of these pre-metastatic niche signatures in involution day 4 samples. We also found the gastric cancer signature was enriched in involution day 6 samples (**Table 1**). Next, we looked at 3 different signatures from MSigDB associated with primary liver cancer. We found none of these hepatocellular carcinoma signatures were enriched during the involution groups. In fact, one signature was significantly downregulated in the involution group (**Table 1**).

**Table 1.** Enrichment scores of liver pathology signatures in involution. Signatures from various liver pathologies compared by GSEA between baseline healthy livers (nulliparous) and involution day 4 or involution day 6 groups. Positive normalized enrichment scores (NES) reflect enrichment in involution samples, whereas negative values reflect enrichment in nulliparous samples.

|  | Pathway | Groups compared | NES | P-val | Reference |
| --- | --- | --- | --- | --- | --- |
| Pre-metastatic niche | COLCAN_SECRETED_PROTEINS_LIVMET | ID4 vs NUL | 2.19 | <0.001 | doi: 10.1186/s13046-023-02833-8 |
|  |  | ID6 vs NUL | 1.61 | 0.011 |  |
|  | PDAC_LIVER_PROMETNICHE | ID4 vs NUL | 1.99 | 0.002 | doi: 10.1038/s41586-019-1004-y |
|  |  | ID6 vs NUL | 1.19 | 0.238 |  |
| Hepatocellular carcinoma | HP_HEPATOCELLULAR_CARCINOMA | ID4 vs NUL | 1.03 | 0.391 | MsigDB: M35273 |
|  |  | ID6 vs NUL | 1.29 | 0.078 |  |
|  | SMITH_LIVER_CANCER | ID4 vs NUL | -1.48 | 0.034 | MsigDB: M18761 |
|  |  | ID6 vs NUL | -1.25 | 0.138 |  |
|  | PATIL_LIVER_CANCER | ID4 vs NUL | -1.31 | 0.064 | MsigDB: M1195 |
|  |  | ID6 vs NUL | -1.23 | 0.118 |  |
| AIH | AIH_HEPATOCYTE_IMMUNE_RESP | ID4 vs NUL | 1.82 | <0.001 | doi.org/10.1101/2023.06.04.23290776 |
|  |  | ID6 vs NUL | 1.47 | 0.020 |  |
|  | AIH | ID4 vs NUL | 1.49 | 0.048 | doi: 10.1371/journal.pone.0264307 |
|  |  | ID6 vs NUL | -1.01 | 0.429 |  |
|  | AIH_CIRRHOSIS | ID4 vs NUL | 0.91 | 0.557 | doi: 10.1371/journal.pone.0264307 |
|  |  | ID6 vs NUL | 0.78 | 0.745 |  |
| Cholestasis | WP_CHOLESTASIS | ID4 vs NUL | -1.00 | 0.462 | MsigDB: M45520 |
|  |  | ID6 vs NUL | -1.00 | 0.431 |  |
|  | HP_CHOLESTATIC_LIVER_DISEASE | ID4 vs NUL | -0.88 | 0.640 | MsigDB: M35918 |
|  |  | ID6 vs NUL | -0.73 | 0.855 |  |
|  | HP_CHOLESTASIS | ID4 vs NUL | -0.83 | 0.923 | MsigDB: M35271 |
|  |  | ID6 vs NUL | 0.93 | 0.708 |  |
| NASH | NASH_VS_NAFL | ID4 vs NUL | -0.97 | 0.476 | doi.org/10.1152/ajpgi.00358.2018 |
|  |  | ID6 vs NUL | -1.24 | 0.167 |  |
|  | NASH_SHH+_VS_NAFL | ID4 vs NUL | 0.91 | 0.619 | doi.org/10.1152/ajpgi.00358.2018 |
|  |  | ID6 vs NUL | -1.30 | 0.095 |  |
|  | NASH_VS_NAFLD+HEALTHYCTRL_UP | ID4 vs NUL | 1.17 | 0.133 | doi.org/10.1038/s41598-022-06512-0 |
|  |  | ID6 vs NUL | 1.31 | 0.058 |  |
| NAFLD | NAFLD_VS_NASH+HEALTHYCTRL | ID4 vs NUL | -1.11 | 0.313 | doi.org/10.1038/s41598-022-06512-0 |
|  |  | ID6 vs NUL | -0.73 | 0.878 |  |
|  | NONALCOHOLIC_FATTY_LIVER_DISEASE | ID4 vs NUL | 1.05 | 0.336 | MsigDB: M39806 |
|  |  | ID6 vs NUL | 1.01 | 0.421 |  |
|  | ADVANCED_NAFLD | ID4 vs NUL | -1.08 | 0.358 | doi.org/10.1126/scitranslmed.aba4448 |
|  |  | ID6 vs NUL | -0.98 | 0.498 |  |
| Cirrhosis | HP_BILIARY_CIRRHOSIS | ID4 vs NUL | 0.91 | 0.598 | MsigDB: M35920 |
|  |  | ID6 vs NUL | 1.10 | 0.320 |  |
|  | HP_CIRRHOSIS | ID4 vs NUL | -0.92 | 0.665 | MsigDB: M35269 |
|  |  | ID6 vs NUL | 0.91 | 0.740 |  |
|  | CIRRHOSIS_POORPROGNOSIS | ID4 vs NUL | 0.92 | 0.623 | doi: 10.1053/j.gastro.2013.01.021 |
|  |  | ID6 vs NUL | 1.12 | 0.216 |  |
| Other | HEPATIC_INSULIN_RESISTANCE_T2DIABETES | ID4 vs NUL | 0.95 | 0.507 | doi: 10.1111/jdi.12998 |
|  |  | ID6 vs NUL | -0.84 | 0.671 |  |
|  | F1_MILD_FIBROSIS_INHEPC | ID4 vs NUL | 1.98 | <0.001 | doi.org/10.1053/j.gastro.2005.09.010 |
|  |  | ID6 vs NUL | 1.84 | <0.001 |  |
|  | OBESE_NAFL_OR_NASH | ID4 vs NUL | -0.82 | 0.714 | doi.org/10.1152/ajpgi.00358.2018 |
|  |  | ID6 vs NUL | -1.03 | 0.430 |  |

Next, we investigated if gene signatures associated with other liver pathologies might be impacted by liver remodeling during involution. We collected signatures from a variety of different liver diseases including: autoimmune hepatitis (AIH), cholestasis, non-alcoholic steatosis (NASH), non-alcoholic fatty liver disease (NAFLD), which has recently been termed metabolic dysfunction-associated steatotic liver disease (MASLD)^26^, cirrhosis, fibrosis, and diabetes. Signatures for AIH were significantly enriched during involution day 4 for two of the three AIH pathways. The AIH gene signature not enriched during involution was an AIH signature associated with cirrhosis, a more advanced liver disease (**Table 1**). We also identified the involution day 4 liver is enriched for a mild fibrosis (F1) signature, but signatures for other more advanced liver pathologies (i.e. NASH, NAFLD, cholestasis, cirrhosis) were not significantly enriched (**Table 1**).

## Discussion

In this study, we investigated previously reported changes in liver remodeling during a lactation-wean cycle, through the exploration of gene expression changes. Previous findings highlight that liver involution has implications in metastatic spread of breast cancer^6,31^, yet liver alterations during a reproductive cycle could have larger implications for liver health and diseases. In our study, we identified lactation as a period of increased liver proliferation and anabolic metabolism. Our findings validate earlier studies describing increased liver size and hepatocyte proliferation during a pregnancy-lactation cycle in rats and mice^6^, and a similar increase in liver size and metabolic output in healthy pregnant women^7^. Previous studies in rodents found that the mammary gland is a site of fatty acid synthesis, especially under fat restricted diets^55^, while others have reported an increase in anabolic enzymes in bovine liver and mammary glands^56^. In addition to increased liver lipid synthesis signatures, we saw an increase in energy producing pathways, including the Krebs cycle. These data concur with previous findings of increased liver NAD metabolites in support of milk production^57^. In sum, these transcriptional profile data are consistent with the need for increased functional output of the liver during lactation. Further, our data strengthen previous studies that defined the functional unit of lactation as including both the mammary gland and the liver^6,58^.

Post-weaning, we identify that the liver shifts to a state of amino acid and lipid catabolism. Transcriptional evidence of these metabolic alterations is apparent as early as involution day 2 and precedes hepatocyte apoptosis and immune infiltrate gene signatures. By day 6 of involution, amino acid catabolism and lipid catabolism pathways decline to pre-pregnancy levels. This suggests that metabolic reprogramming to a baseline state is largely completed by 6 days post-wean. Our observation that the shift to catabolic metabolism precedes liver tissue remodeling (i.e. apoptotic and immune cell influx signatures) emphasizes the importance of metabolic demand on liver size and function. Interestingly, studies in mice have shown that pups can be given back to the dam within two days of mammary gland involution and milk production will be re-established, but not after 2 days post-wean^59,60^. Given that catabolism rapidly increases in the liver following day 2, we suspect triggering of catabolic pathways in the liver may be a mechanism that prevents reestablishment of milk production in mammary gland.

We identified increases in apoptotic gene signatures during weaning-induced liver involution after the shift to a catabolic state, but not necrosis or autophagic cell death. Apoptosis is classically described as immune silent because mechanisms, such as efferocytosis of dying cells^13^, minimize destructive inflammatory responses^61^. Apoptosis is the primary mechanism of cell death during embryonic development^62^, thus our identification of apoptosis as the main mechanism of cell death during weaning-induced involution is consistent with a programmed, immunosuppressive mechanism. Further analyses of liver involution revealed an apoptotic pathway associated with lysosomal content leakage. Similar observations have been made in the mammary gland, whereby milk stasis leads to endocytosis of fat globules by mammary epithelial cells^41,42^. These fat globules, broken down into free fatty acids in the lysosome, have been shown to permeabilize lysosomal membranes, resulting in the release of proteases such as cathepsins, and STAT3 mediated signaling^42^ resulting in lysosomal mediated cell death^41,44^. Here, we provide correlative evidence in the involuting liver. We show an increase in lipid uptake, followed by a release of proteases from the lysosome, which corresponds with peak apoptosis signals during liver involution. However, further mechanistic studies are needed to show a causal relationship between post-wean lipid abundance, increased lysosomal leakiness, and hepatocyte cell death.

Our data on potential mechanisms of cell death add to our current understanding of weaning-induced liver involution, and further supports functional interdependence between the liver and the mammary gland. Relevance to women is implicated by similar liver size increase with pregnancy and return to baseline post-wean^7^. Further, in women, lack of liver gain-loss with a pregnancy cycle was found to associate with increased gestational hypertension and reduced insulin sensitivity^7^. Combined these data support the concept that healthy pregnancy in rodents and women includes a gain in liver size and post-wean loss, emphasizing liver involution as a physiologically normal and pre-programmed feature of a reproductive cycle.

Along with the identification of apoptosis signatures during involution, we also identified evidence suggesting the involuting liver is immune suppressed. Here, we show transcriptomic and histologic evidence for increased lipid filled macrophages during involution, which have been previously associated with immune suppression^52,63^. One study reported that enhanced fatty acid metabolism can skew macrophage polarization to an immune suppressive macrophage^63^, while another demonstrates STAB1+TREM2^high^ lipid filled macrophages could inhibit T-cell activation and proliferation^52^. While we do not demonstrate mechanism, we theorize that lipids shuttled from the mammary gland to the liver are taken up not only by hepatocytes, but also macrophages, and that lipid filled macrophages contribute to suppression of immune activation during the cell death phase of liver involution. In addition to identifying an increase in lipid filled macrophages, we also identify an increase in CD45+CD11b+Gr1+ myeloid cells. While this phenotype is shared with normal monocytes, it has also been used to identify mouse MDSCs^47^, which are classically associated with immune suppression. We postulate that since liver involution is a normal facet of reproduction, mechanisms to prevent involution from triggering a more tissue damaging inflammatory response would be in place. The presence of increased lipid loaded macrophages and MDSCs during involution are consistent with prior studies demonstrating reduced T cell response to antigen stimulation in the liver during involution^8^. However, a limitation to our study is we do not functionally test the immune suppressive capabilities of liver lipid filled macrophages nor immature myeloid cells associated with involution.

The large shifts in the liver transcriptome across a reproductive cycle also correlate with some liver pathologies. We found that liver pre-metastatic niche signatures were enriched during the involution period. These data support previous findings showing increased risk of breast cancer liver metastasis in rodent models of postpartum breast cancer (PPBC) and in women diagnosed with PPBC^6,8,31^. Interestingly, the pre-metastatic niche signatures we identified were generated from different primary cancers (i.e. pancreatic, colorectal, and gastric), suggesting that liver involution may generate a pre-metastatic niche that is broadly favorable across cancer types. While this is an untested hypothesis, it is supported by clinical case studies which identified progression to liver metastases from various primary cancers (i.e. pancreatic, rectal, and ovarian) in patients after a recent childbirth^64^.

Besides recent studies in PPBC and liver metastasis, the relationship between parity status and liver pathologies has been sparsely studied to date. To our surprise, the involuting liver does express gene signatures consistent autoimmune hepatitis (AIH), but does not transcriptionally resemble many of these extreme disease-state livers (i.e. cholestasis, NASH, NAFLD, cirrhosis). Interestingly, several studies have shown an increase in severity of AIH during the postpartum period^65,66^, with one study reporting 20-50% of patients with AIH report flare ups during the first several months postpartum^67^. Another case study traced the liver dysfunction of three postpartum women to AIH within several months after giving birth^68^. We hypothesize that these postpartum AIH flare ups could be exacerbated by inflammation occurring during liver involution. However, these studies on AIH postpartum do not report weaning times. Many studies are done with relation to birth time but given our findings that emphasize the post-wean involution period as a dramatic window of tissue remodeling, we recommend clinicians consider reporting weaning time.

In sum, our study demonstrates complex and transient functional states of the liver over a lactation-wean cycle. These findings underscore the liver’s adaptive metabolic and immune responses needed to accommodate the metabolic demands of lactation and weaning. This study also has potential implications for liver pathologies, such as increased risk of metastasis and AIH. Further, this dataset can serve as a resource for others to better understand how liver biology might impact maternal and newborn health, and possibly contribute to sex-specific differences in liver pathologies.

## Methods

### Sample Description

Archival snap frozen liver tissue from BALB/C mice at different reproductive timepoints was used as starting material. Reproductive groups were generated using methods previously reported^20^, nulliparous (Nul) (n=4), day 10 of lactation (Lac) (n=3), involution day 2 (ID2) (n=3), ID4 (n=4), ID6 (n=6), ID8 (n=3), and regression (n=4). Samples were taken from three independent studies. Mice for these three studies were ordered from Charles River Laboratories and Jackson Laboratories. Ethics approval was obtained from the Oregon Health & Science University (Approval number TR02_IP00000967).

### RNA Isolation

Liver was snap frozen in LN2, and ground into a fine powder by mortar and pestle. RNA was extracted using trizol and the Fischer Scientific Zymo Direct-zol RNA MiniPrep kit (50-197-7077). RNA quality and concentration was assessed by the Gene Profiling Shared Resource Core at Oregon Health and Science University, using the Aglient 2100 BioAnalyzer.

### Library Preparation and Sequencing

Library preparation and sequencing was completed by Novogene. Novogene used NEBNext® Ultra RNA Library Prep Kit for Illumina® (cat# E7420S, New England Biolabs, Ipswich, MA, USA) according to the manufacturer’s protocol. This resulted in 250-350 bp insert libraries which were quantified using the Qubit 2.0 fluorometer (Thermo Fisher Scientific, Waltham, MA, USA) and quantitative PCR. Size distribution was analyzed using an Agilent 2100 Bioanalyzer (Agilent Technologies, Santa Clara, CA, USA). Qualified libraries were sequenced on an Illumina Novaseq6000 Platform (Illumina, San Diego, CA, USA) using a paired-end 150 run (2×150 bases). The library input was standardized for 100ng of input cDNA.

### RNA Sequence Alignment, Data Processing, and Analysis

The raw fastq files were first quality checked using FastQC (version 0.11.8). Fastq files were aligned to mm10 mouse reference genome (GRCm38.89) via STAR (version 2.6.1a) and per-gene counts quantified by RSEM (version 1.3.1) based on the gene annotation Mus_musculus.GRCm38.89.chr.gtf. Differential gene expression values were obtained using DESeq2 (version 1.42.1). Gene expression differences were considered significant if they passed the following criteria: adjusted P value < 0.05, log2(fold change) ≥ 1.5. Raw read counts were processed and normalized to transcripts per million (TPM) using the RSEM algorithm (v1.3.1). The sequencing data has been deposited in the National Center for Biotechnology Information/Gene Expression Omnibus under accession number GSE260714.

### STRING & PANTHER Analysis

STRING analysis (https://string-db.org/) was run on the top 500 most significant differentially expressed genes (DEGs), with both up and down genes included. K-mean clustering was done to identify the main clusters of related DEGs. Genes included within each cluster were put into PANTHER GO Enrichment Analysis (https://geneontology.org/) to reduce related GO terms into related families of GO terms, being listed under a “parent” GO term. The top 5 most significant parent GO terms were plotted using GraphPad Prism.

### Gene Set Enrichment Analysis (GSEA)

Gene set enrichment analysis (GSEA) v 4.2.3^69^ was used to identify pathways enriched between different reproductive time points. We used the and the chip platform Mouse_ENSEMBL_Gene_ID_Human_Orthologs v7.5.chi. Signatures were run using 1000 gene set permutations. Enrichment statistics were weighted, and ranked by signal to noise. We used Deseq2 normalized values. We ran gene sets from MSigDB collections^37^ including: Hallmark, Biocarta, Kegg, and Reactome (**Supplementary Table 2**). We also ran GSEA on customized gene sets from various studies^47,48,50,53,54^ (**Table 1**). Pathways were considered enriched if they had a normalized p-value <0.05.

### Single sample Gene Set Enrichment Analysis (ssGSEA)

Single sample GSEA (ssGSEA)^36^ was used to identify enrichment of pathways between all groups. ssGSEA was run using GenePattern^70^ using the same signatures used in our GSEA analysis. We used log converted transcript per million (TPM) reads as our input for these analyses, with no additional normalization methods, a weighting exponent of 0.75, and using gene sets no smaller than 10. Individual ssGSEA plots were plotted using GraphPad Prism, using multiple comparisons, one-way ANOVA for statistical measurements. Enrichment scores were plotted using GraphPad Prism and one-ANOVA tests were run. Heat maps were generated using Morpheus(https://software.broadinstitute.org/morpheus).

### Regulon Analysis

To infer transcription factor activity, we used the master regulator inference algorithm (MARINa) compiled in R ‘viper’ package to perform the regulon analyses on nulliparous, lactation, and involution liver samples. In each comparison, two source gene expression signatures were used and regulatory networks were required as model inputs. We used Student’s t-test based statistics, as suggested in the viper manual, was used. The regulons used for the transcription factor activity inference were curated from four databases. The single sample-based regulon activities were inferred by function “viper”, which is an extension of MARINa114 and transforms a gene expression matrix to a regulatory protein activity matrix. For the model input, we used the TPM quantification of different reproductive stage liver samples as the expression matrix and the same regulon network described above as the regulatory network.

### Immunohistochemistry (IHC), multiple immunohistochemistry (mIHC), and immunofluorescence (IF)

Tissue samples were fixed in 10% NBF for 48 hours with shaking, at which point the tissue was placed in 70% ethanol until processing. Tissue was then processed in a Sakura Tissue-Tek VIP 5 tissue processor for 11 hours. Samples were embedded in paraffin wax. Tissues were sectioned into 4μm sections on ionized glass slides (Mercedes Scientific, TNR WHT90ADCC) and allowed to dry at 37°C for at least 24 hours. When ready to stain, slides were deparaffinized at 60C for 1 hour. Slides were washed in xylene 4 times for 5 minutes each for a total of 20 minutes to remove paraffin. Slides were then rehydrated using several quick washes in 100%, then 95%, then 70% ethanol, then 50% ethanol, followed by 2 washes in de-ionized water. Slides were placed into 1x TBST for immediate staining or stored in 1x TBS at 4°C for up to 48 hours before staining. Antigen retrieval was done using 1x EDTA diluted from a 5x stock solution (pH 9.0) (Biocare Medical, CB917M) at 115°C for 20 minutes in a pressure cooker (BioSB, BSB 7087). Tissue was blocked with 1% Triton X-100 (Sigma Aldritch, X100) + 5% Normal Goat Serum (Sigma Aldrich, S26-M) + 2.5% Bovine Serum Albumin (Fisher Scientific BP1600-100) in PBS for 30 minutes. Tissue that was stained for IHC, IF, or mIHC was blocked for an additional 15 minutes in 30% hydrogen peroxide diluted in methanol for a 3% hydrogen peroxide solution. Tissue fixation and processing was validated by staining all tissues for E-cadherin expression, and all tissues passed quality control. For experimental IHC and mIHC we stained the following antibodies for 1 hour at room temperature at the described concentrations: ADPH (1:1000, Novus Biologicals Cat: NB110-40878), CD45 (1:50, BDPharminigen, Cat: 550539), Gr1 (1:100, Invitrogen, Cat: 14-5931-82). Secondary antibodies conjugated with horseradish peroxidase were diluted at 1:200, and stained for 30 minutes. Tissue was counterstained with hematoxylin for 1 minute at room temperature (Agilent Technologies-Dako, Cat S330130-2). For IF, slides were prepared in the same method as described above. The following antibodies were stained for 1 hour at room temperature at the described concentrations: F4/80 (1:100, Thermo Fisher Cat: MA1-91124 70076S), ADPH (1:1000, Novus Biologicals Cat: NB110-40878), Ctsb (1:2000, Cell Signaling, Cat: 31718T). Lamp1 was stained overnight (1:1000, Cell Signaling, Cat: 9091). Secondaries were applied at for 30 minutes at room temperature, protected from light: Rabbit-AF 647 (1:250, Invitrogen, A-21245), Rat-AF488 (1:250, Invitrogen, A11006). Slides were mounted using TrueVIEW® Autofluorescence Quenching Kit (Vector Laboratories, SP-8400-15) and stored at 4°C until ready for imaging. IHC and mIHC slides were scanned on the Aperio AT2 slide scanner (Leica Biosystems, Wetzlar, Germany). Stained area analyses were done using Aperio Image Scope digital pathology software (version 12.4.6.5003). For adipophilin quantification, 75µm circles were drawn around portal triads or central veins, and percent area was quantified for 3 portal triads and 3 central veins was done per mouse. For mIHC data analysis, slides were analyzed first using MatLab (version R2022a) for registration of slides and alignment of stains, then taken into CellProfiler (CellProfiler version 4.2.6) for creating masking images. Finally, using FCS Express software (version 7.18.0025) slides were gated into different groups based on whether a cell was stained positively or negatively for an antibody. For immunofluorescence studies, slides were imaged using the Zeiss Axio Observer microscope and ApoTome 2 imaging system at 20x. Analysis was done using Zeiss ZEN Blue software from OHSU’s Advanced Light Microscopy Core.

## Notes

### Competing Interest Statement

The authors have declared no competing interest.

