## Supplementary Tables and Figures for "Reproductive state controls transcription in the murine liver, with implications for breast cancer liver metastasis"

**Supplementary Table 1.** Different cell death mechanisms were evaluated by GSEA. Normalized enrichment scores (NES) and nominal p-values were recorded for involution day 4 livers as compared to lactation livers. Positive NES values reflect enrichment in involution samples, whereas negative values reflect enrichment in lactation samples. Statistically significant pathways are shown in white, and non-statistically significant values are shown in grey.

| | **Pathway** | **NES** | **NOM P-val** | **MSigDB Ref** |
|---|---|---|---|---|
| Apoptosis | GOBP_EXTRINSIC_APOPTOTIC_SIGNALING_PATHWAY | 1.61 | 0.006 | MsigDB: M11317 |
| | GOBP_REGULATION_OF_EXTRINSIC_APOPTOTIC_SIGNALING | 1.48 | 0.002 | MsigDB: M13195 |
| | WP_APOPTOSIS | 1.42 | 0.024 | MsigDB: M39664 |
| | HALLMARK_APOPTOSIS | 1.39 | 0.012 | MsigDB: M5902 |
| | KEGG_APOPTOSIS | 1.34 | 0.055 | MsigDB: M8492 |
| Non-Apoptotic Pathways | REACTOME_REGULATED_NECROSIS | 1.36 | 0.065 | MsigDB: M41803 |
| | GOBP_NECROTIC_CELL_DEATH | 1.26 | 0.117 | MsigDB: M24757 |
| | HP_HEPATIC_NECROSIS | 0.68 | 0.853 | MsigDB: M35916 |
| | BIOCARTA_DEATH_PATHWAY | 0.91 | 0.589 | MsigDB: M14971 |
| | GOBP_AUTOPHAGIC_CELL_DEATH | 1.06 | 0.392 | MsigDB: M23831 |

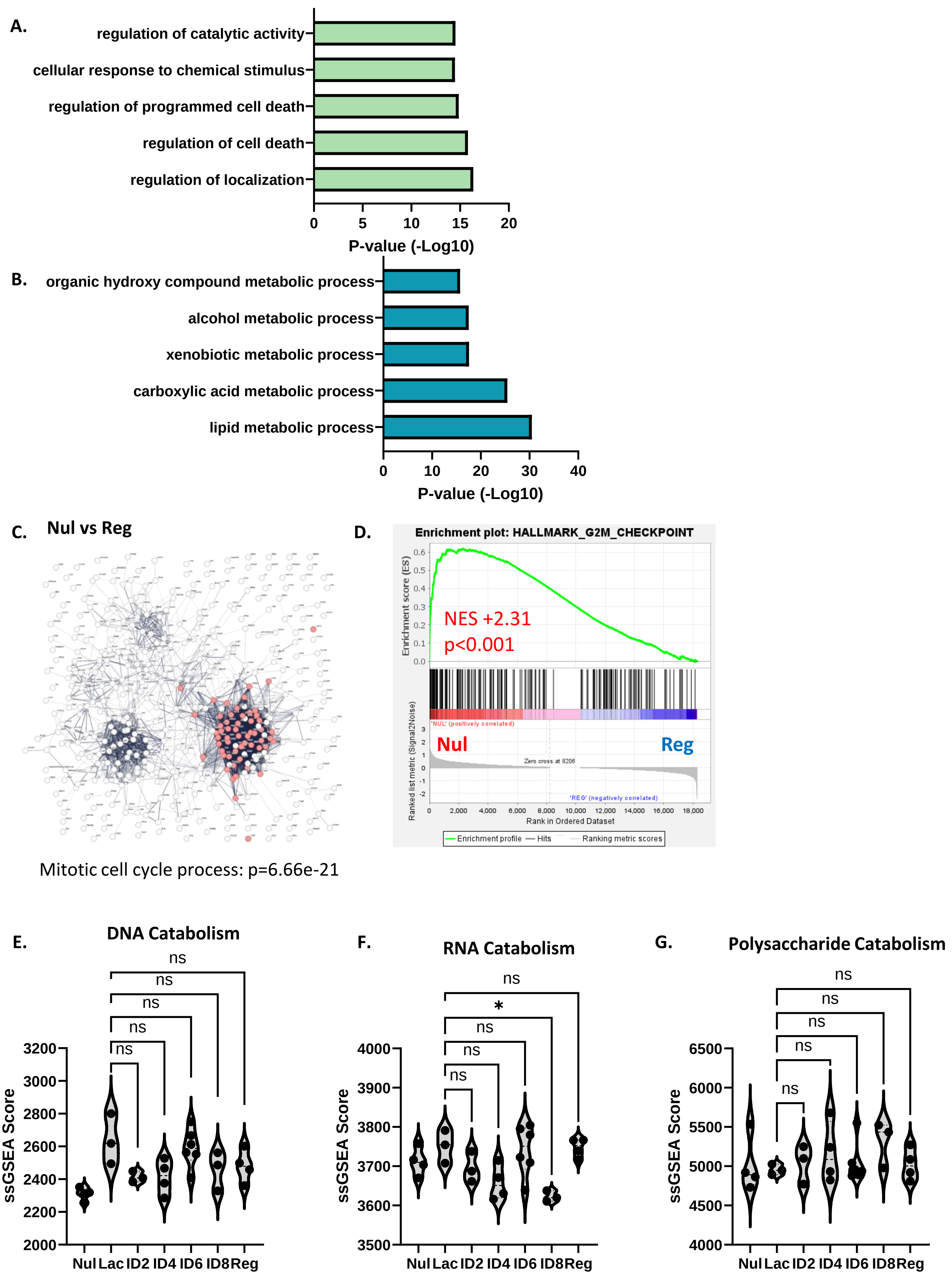
A.
regulation of catalytic activity
cellular response to chemical stimulus
regulation of programmed cell death
regulation of cell death
regulation of localization
0
5
10
15
20
P-value (-Log10)
B.
organic hydroxy compound metabolic process
alcohol metabolic process
xenobiotic metabolic process
carboxylic acid metabolic process
lipid metabolic process
0
10
20
30
40
P-value (-Log10)
C.
Nul vs Reg
Mitotic cell cycle process: p=6.66e-21
D.
Enrichment plot: HALLMARK_G2M_CHECKPOINT
Enrichment score (ES)
NES +2.31
p<0.001
'NUL' (positively correlated)
Nul
Reg
Zero cross at 8206
'REG' (negatively correlated)
Ranked list metric (Signal2Noise)
Rank in Ordered Dataset
Enrichment profile
Hits
Ranking metric scores
E.
DNA Catabolism
ns
ns
ns
ns
ns
ssGSEA Score
Nul Lac ID2 ID4 ID6 ID8 Reg
F.
RNA Catabolism
ns
*
ns
ns
ns
ssGSEA Score
Nul Lac ID2 ID4 ID6 ID8 Reg
G.
Polysaccharide Catabolism
ns
ns
ns
ns
ns
ssGSEA Score
Nul Lac ID2 ID4 ID6 ID8 Reg

**Supplementary Fig 1.** GO Term Enrichment analysis of the top 5 most enriched pathways in nulliparous compared to lactation livers within the **A)** green cluster and the **B)** blue STRING clusters. **C)** STRING analysis between nulliparous and regression groups shows enrichment for cell cycle regulated processes. **D)** GSEA analysis of the Hallmark G2M pathway between nulliparous and regression livers. **E)** ssGSEA analysis of catabolism of DNA nucleotides, **F)** mRNA, **G)** and polysaccharides (one-way ANOVA, mean comparisons to lactation, *$p<0.05$).

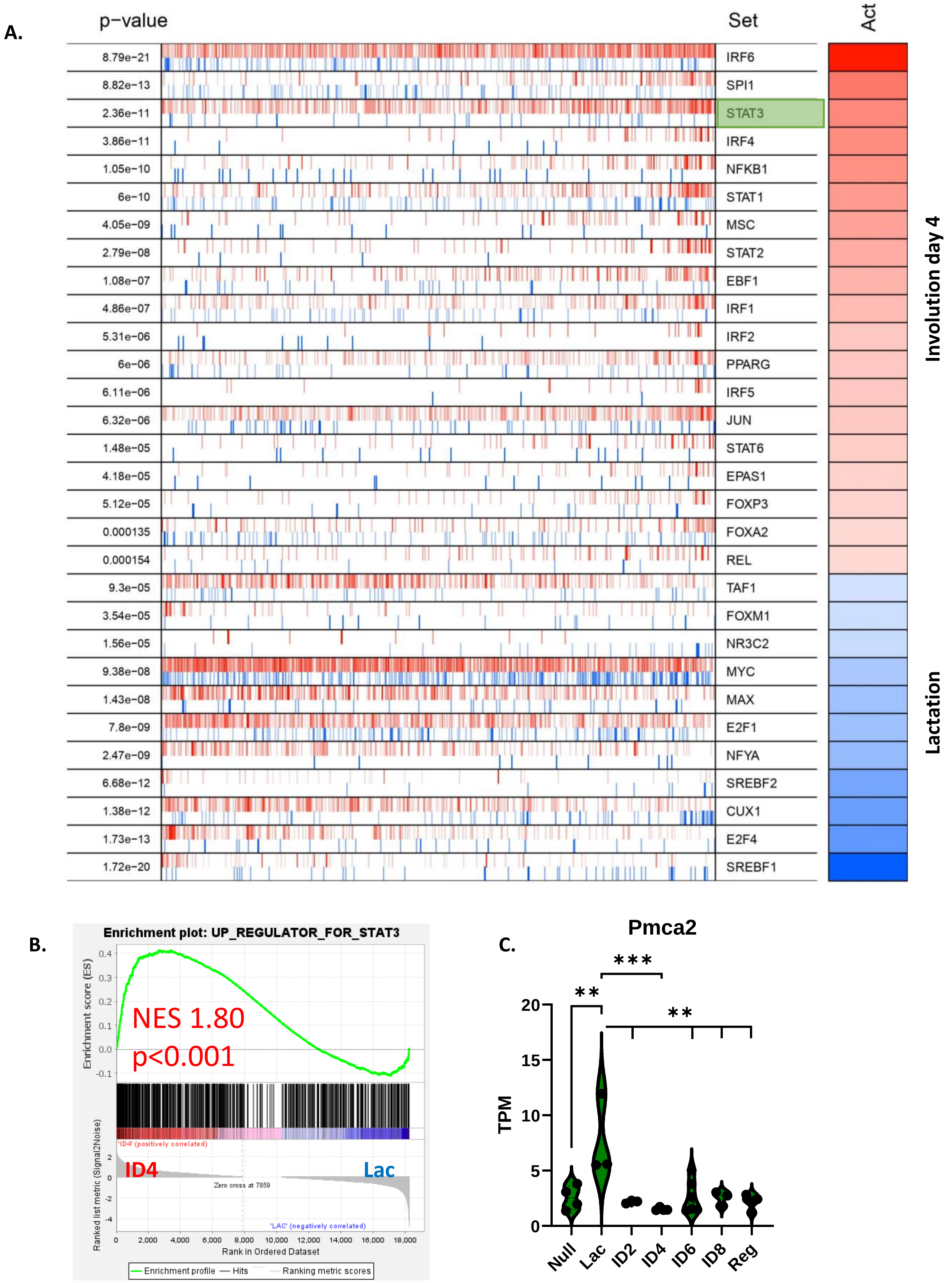


**Supplementary Fig 2. A)** Regulon analysis between lactation (blue) and involution day 4 (red) groups. **B)** Enrichment of STAT3 pathway genes compared between lactation (blue) and involution day 4 (red) livers by GSEA. **C)** Gene expression of Pmca2 and reported as transcripts per million (TPM).

**A.** CibersortX LM22

| Mixture | B cells naive | B cells memory | Plasma cells | T cells CD8 | T cells CD4 naive | T cells CD4 memory resting | T cells CD4 memory activated | T cells follicular helper | T cells regulatory (Tregs) | T cells gamma delta | NK cells resting | NK cells activated | Monocytes | Macrophages M0 | Macrophages M1 | Macrophages M2 | Dendritic cells resting | Dendritic cells activated | Mast cells resting | Mast cells activated | Eosinophils | Neutrophils |
|---|---|---|---|---|---|---|---|---|---|---|---|---|---|---|---|---|---|---|---|---|---|---|
| NUL_227_TPM | 0.01619895 | 0.07780537 | 0.31762737 | 0.0407442 | 0.05884344 | 0 | 0.01183721 | 0 | 0.01197894 | 0 | 0.02785809 | 0.03297528 | 1.03712802 | 0.07192977 | 0.02366145 | 0 | 0.12051229 | 0.01502877 | 0.09537219 | 0 | 0 | 0 |
| NUL_226_TPM | 0 | 0.13279327 | 0.34280196 | 0.02662143 | 0.04527176 | 0 | 0 | 0.03637864 | 0 | 0 | 0.09942985 | 0 | 0.97709086 | 0.04191656 | 0.0714139 | 0.01901956 | 0.11094267 | 0 | 0.08623084 | 0 | 0.01355071 | 0 |
| NUL_220_TPM | 0 | 0.13908565 | 0.188923 | 0.05093754 | 0 | 0.06672292 | 0 | 0.02982478 | 0.01405642 | 0 | 0.04299943 | 0.05801122 | 1.03151029 | 0.06740897 | 0.06810512 | 0.00540874 | 0.09201321 | 0 | 0.09018936 | 0 | 0.00373535 | 0 |
| NUL_221_TPM | 0 | 0.23942695 | 0.19994264 | 0.02879906 | 0.01598734 | 0.01401143 | 0.01851544 | 0.00205593 | 0.02779839 | 0 | 0.04695115 | 0.01807945 | 0.96182802 | 0.12822007 | 0.03807517 | 0 | 0.08077663 | 0 | 0.07825936 | 0 | 0 | 0 |
| LAC_200_TPM | 0.00838387 | 0.09485007 | 0.36067601 | 0.0019817 | 0.02401444 | 0 | 0.00779419 | 0.04363665 | 0 | 0 | 0.02611692 | 0.03666293 | 0.82367871 | 0.07705781 | 0.03545654 | 0 | 0.14567313 | 0 | 0.09295547 | 0 | 0.01007329 | 0 |
| LAC_240_TPM | 0.02466217 | 0.0970598 | 0.41705175 | 0 | 0.04827206 | 0.01373816 | 0 | 0.01409271 | 0 | 0 | 0.08343207 | 0.00264876 | 0.81519575 | 0.04876629 | 0.03306932 | 0 | 0.0915385 | 0 | 0.04584494 | 0 | 0.01103777 | 0 |
| LAC_213_TPM | 0.01420942 | 0.06954547 | 0.2780711 | 0 | 0 | 0.0520242 | 0 | 0.01689092 | 0.00823487 | 0 | 0.02804526 | 0.03807747 | 0.83778564 | 0.05183274 | 0.03900306 | 0 | 0.13668259 | 0 | 0.10354952 | 0 | 0 | 0 |
| ID2_39_TPM | 0.00737815 | 0.10583485 | 0.28643558 | 0.02548652 | 0.02280476 | 0.00193062 | 0 | 0.01517608 | 0 | 0 | 0.05005114 | 0.01031782 | 0.98633371 | 0.06860306 | 0.05566445 | 0 | 0.16842876 | 0 | 0.06642884 | 0 | 0.01072133 | 0 |
| ID2_31_TPM | 0.01427849 | 0.141014 | 0.34036659 | 0.07054997 | 0 | 0.01599951 | 0 | 0.05204072 | 0.00517147 | 0 | 0.00367659 | 0.05245221 | 0.93672666 | 0.09227497 | 0.05403637 | 0 | 0.11986352 | 0 | 0.08814349 | 0 | 0 | 0 |
| ID2_224_TPM | 0.0057023 | 0.09498457 | 0.28321036 | 0.03284378 | 0 | 0.05571185 | 0.01688942 | 0.00945546 | 0 | 0 | 0.04488592 | 0.01199035 | 0.94605288 | 0.06061706 | 0.0582975 | 0 | 0.18477918 | 0 | 0.08551734 | 0 | 0.00933274 | 0 |
| ID4_215_TPM | 0 | 0.16441124 | 0.2633883 | 0.05019567 | 0.0373387 | 0 | 0 | 0 | 0.00535246 | 0.01487795 | 0.05705164 | 0 | 1.08275883 | 0.09446949 | 0.05703729 | 0 | 0.16542278 | 0 | 0.04594363 | 0 | 0 | 0 |
| ID4_211_TPM | 0 | 0.20818369 | 0.21006758 | 0.02792584 | 0 | 0.02083607 | 0.03592726 | 0 | 0.01011368 | 0.00612289 | 0.03937215 | 0.00814901 | 1.12211791 | 0.07409899 | 0.05921762 | 0 | 0.12445488 | 0 | 0.08407825 | 0 | 0 | 0 |
| ID4_13_TPM | 0.00285041 | 0.10851889 | 0.27873328 | 0.01353988 | 0.03845441 | 0 | 0 | 0.01841399 | 0 | 0 | 0.05750049 | 0.05287215 | 1.19208618 | 0.06924349 | 0.05667021 | 0.00797115 | 0.11288855 | 0 | 0.08083195 | 0 | 0.00206196 | 0 |
| ID4_15_TPM | 0.00196593 | 0.10245631 | 0.31509767 | 0.05470041 | 0.00342085 | 0 | 0.00158458 | 0.04846201 | 0.002749 | 0 | 0.05637037 | 0.0161551 | 1.09383432 | 0.05742424 | 0.08026149 | 0 | 0.22655205 | 0 | 0.07659379 | 0 | 0.02809246 | 0 |
| ID6_218_TPM | 0 | 0.17559995 | 0.31953511 | 0.05818979 | 0.01373563 | 0 | 0.02249763 | 0.00489468 | 0.02713713 | 0 | 0.07531493 | 0.01462304 | 1.18120197 | 0.13128065 | 0.06327379 | 0 | 0.10770643 | 0 | 0.08970808 | 0 | 0 | 0 |
| ID6_201_TPM | 0 | 0.13526736 | 0.22925553 | 0.04391468 | 0.05901408 | 0 | 0 | 0 | 0.05195897 | 0.00322065 | 0.05712502 | 0.0164197 | 1.17053011 | 0.09577952 | 0.04416984 | 0 | 0.11366802 | 0 | 0.10193101 | 0 | 0 | 0 |
| ID6_216_TPM | 0.01045456 | 0.0763736 | 0.39069319 | 0.04490583 | 0.01765367 | 0 | 0 | 0.06090364 | 0.00166286 | 0 | 0.03981395 | 0.0155758 | 1.1324193 | 0.10813378 | 0.04901828 | 0 | 0.11887422 | 0 | 0.07598586 | 0 | 0 | 0 |
| ID6_209_TPM | 0 | 0.12032642 | 0.31430947 | 0.04230974 | 0.0221952 | 0 | 0 | 0.02883378 | 0.06054299 | 0 | 0.01596482 | 0.04350032 | 1.07074211 | 0.09016112 | 0.03521369 | 0 | 0.14919907 | 0 | 0.1036694 | 0 | 0 | 0 |
| ID6_232_TPM | 0 | 0.15359376 | 0.29672924 | 0.05637765 | 0.00755519 | 0 | 0 | 0.04503212 | 0.00013148 | 0.00033634 | 0.11759159 | 0 | 1.22189953 | 0.06929499 | 0.06757775 | 0 | 0.11938052 | 0 | 0.03348232 | 0 | 0 | 0 |
| ID6_230_TPM | 0.00780227 | 0.16098661 | 0.29673848 | 0.0259237 | 0.02723098 | 0.02675864 | 0.04393749 | 0 | 0.02742376 | 0 | 0.10865546 | 0 | 1.04353706 | 0.11592321 | 0.06685758 | 0 | 0.15857076 | 0 | 0.06822726 | 0 | 0 | 0 |
| ID8_98b_TPM | 0.0134157 | 0.18464912 | 0.27559927 | 0.0300481 | 0.03949158 | 0 | 0.00370979 | 0 | 0.03542523 | 0.01142687 | 0.07756436 | 0.00362346 | 0.96782104 | 0.12234773 | 0.06177749 | 0 | 0.12532879 | 0 | 0.08571623 | 0 | 0 | 0 |
| ID8_101b_TPM | 0.00516486 | 0.12644782 | 0.2810506 | 0.01885286 | 0.03762862 | 0.0301756 | 7.2512E-05 | 0.00549225 | 0.00861882 | 0.01918901 | 0.02473197 | 0.03750399 | 0.92689593 | 0.06051092 | 0.04133515 | 0.03619659 | 0.13503098 | 0 | 0.1235883 | 0 | 0.00378526 | 0 |
| ID8_14_TPM | 0.01283338 | 0.12373925 | 0.35675716 | 0.03399559 | 0.02205814 | 0 | 0 | 0.02780262 | 0.01033763 | 0 | 0.06241764 | 0.02445149 | 1.01803256 | 0.05958908 | 0.04673697 | 0.03016365 | 0.13370986 | 0 | 0.11727308 | 0 | 0.00848857 | 0 |
| REG_239_TPM | 0 | 0.23047527 | 0.2761999 | 0.12815027 | 0 | 0 | 0 | 0.02183708 | 0.04818817 | 0 | 0.07265407 | 0.03030758 | 0.92328945 | 0.09656756 | 0.05142545 | 0 | 0.14206423 | 0 | 0.05180531 | 0 | 0 | 0 |
| REG_214_TPM | 0 | 0.16998481 | 0.32173846 | 0.05667022 | 0.00702586 | 0.02040799 | 0.06819655 | 0 | 0.0220881 | 0 | 0.08281444 | 0 | 1.01001427 | 0.07141792 | 0.06021596 | 0 | 0.10609091 | 0 | 0.04097948 | 0 | 0 | 0 |
| REG_234_TPM | 0.01279531 | 0.14555912 | 0.32988301 | 0.03745963 | 0 | 0.05534589 | 0.04344084 | 0 | 0.0434973 | 0 | 0.08670409 | 0 | 0.96885488 | 0.09353959 | 0.03720122 | 0 | 0.20008844 | 0 | 0.04110476 | 0.00027057 | 0 | 0 |
| REG_203_TPM | 0 | 0.16664557 | 0.27611733 | 0.07017194 | 0 | 0.06050085 | 0.01268807 | 0.0012395 | 0.01472117 | 0 | 0.01955097 | 0.03610854 | 1.0861013 | 0.13271434 | 0.06937909 | 0 | 0.10573079 | 0 | 0.08916851 | 0 | 0 | 0 |

**B.** CD45+ CD11b+ Ly6C/G+ Myeloids

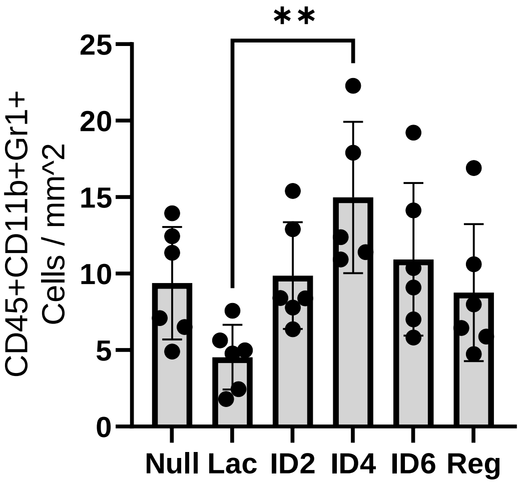


**C.** **Neutrophil Specific**

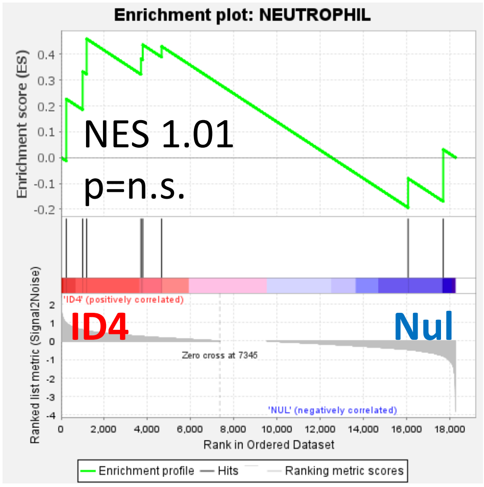


**D.** **Cibersort: M1 Macrophages**

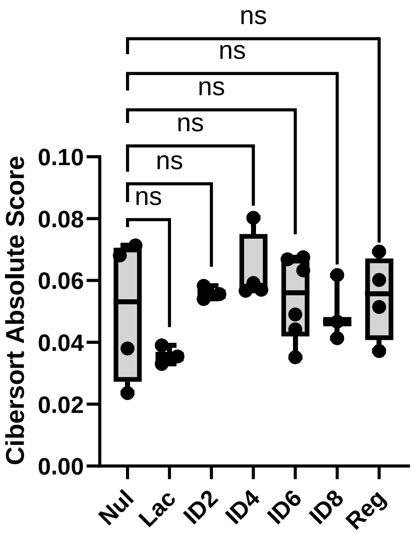


**E.** **Null vs ID4** **Null vs ID6**

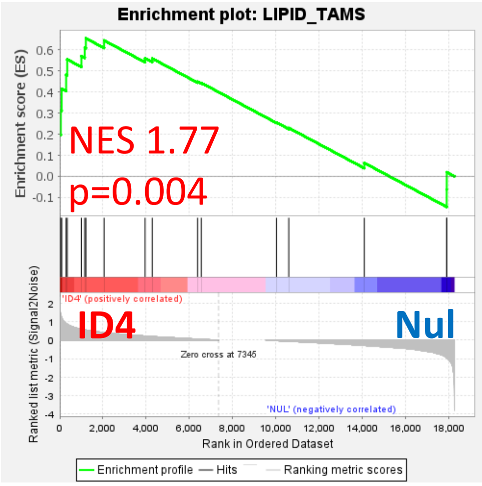


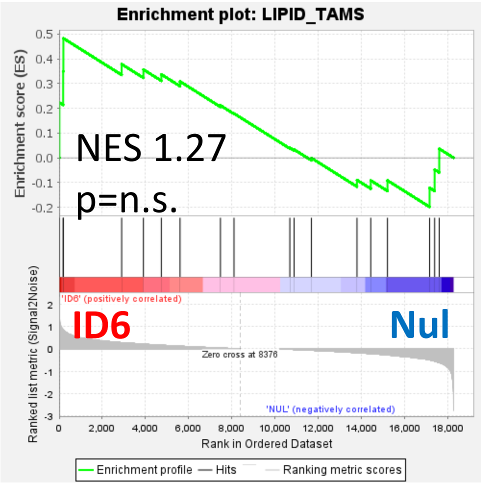


**Supplementary Figure 3. A)** CibersortX values using the LM22 signature to identify different immune cell subsets that were upregulated (red) or downregulated (blue) across reproductive time points. **B)** Quantification of mIHC CD45+CD11b+Gr1+ myeloid cells per mm^2 area. Average of 4 regions of analysis per mouse liver (n=5-6 livers/group, one-way ANOVA, **p<0.01). **C)** GSEA analysis on a signature specific to neutrophils between nulliparous and involution day 4 groups. **D)** CibersortX analysis on the LM22 M1 macrophage signature (one-way ANOVA). **E)** GSEA analysis on a signature for lipid associated macrophages between nulliparous and involution day 4 livers (left) or involution day 6 (right).
